# Purposeful Design of a Survey Tool to Evaluate the Adequacy of Hospitals’ Water Sanitation and Hygiene and Allocate Responsibility for Action – from Wash FIT to Wash Fast

**DOI:** 10.1101/622449

**Authors:** Michuki Maina, Mathias Zosi, Grace Kimemia, Paul Mwaniki, Arabella Hayter, Margaret Montgomery, Jacob McKnight, Olga Tosas-Auguet, Constance Schultsz, Mike English

## Abstract

**Background:** Poor water sanitation and hygiene (WASH) in health care facilities increases hospital associated infections and results in greater use of second line antibiotics, which drives antimicrobial resistance. The existing assessment tool, Water and Sanitation for Health Facility Improvement Tool (WASH FIT), is designed for self-assessment in smaller primary facilities. A tool is needed for larger facilities with multiple inpatient units, that supports comparison of multiple facilities and identifies who is responsible for action at different levels of the health system.

**Methods:** We adapted the WASH FIT tool to: 1) create a simple numeric scoring approach to enable comparison of hospitals and facilitate tracking of WASH performance over time; (2) identify indicators that can be assessed and scored for each hospital ward to help identify variation within facilities and; (3) identify those responsible to effect positive change at different levels of the health system. We used a pilot, analysis of interview data and consultative stakeholder meetings to establish the feasibility and face validity of the WASH Facility Survey Tool (WASH FAST).

**Results:** WASH FAST can be used to produce an aggregate percentage score at facility level to summarise hospitals’ overall WASH status and illustrate variation across hospitals. Thirty-four of the 65 indicators spanning four WASH domains can be assessed at ward level enabling between ward variations to be highlighted. Three levels of responsibility for WASH service monitoring and improvement were identified that were supported by qualitative data and multiple stakeholders: the county/regional level, hospital senior management and the infection prevention and control committee within the healthcare facility.

**Conclusion:** We propose WASH FAST can be used as a survey tool to assess, improve and monitor progress of WASH and IPC in hospitals in resource-limited settings, providing useful data for decision making and contributing to wider quality improvement efforts.

## BACKGROUND

Improving water supply, hygiene, sanitation and healthcare waste management (WASH) is captured under Sustainable Development Goal 6 (1). Improving WASH in healthcare facilities (HCFs) is linked to specific benefits, including reductions in hospital associated infections and antimicrobial resistance; better management and control of disease outbreaks, improved staff morale and a reduction in healthcare costs (2) (3). It also influences communities – as health staff model proper hygiene practices (4) – and may improve patients’ trust in and experience of care and subsequently their satisfaction with and uptake of health services (5) (6).

Recent data released by the World Health Organization(WHO)/ United Nations children’s Fund (UNICEF) Joint Monitoring Programme for Water Supply, Sanitation and Hygiene (JMP) suggests that the health facilities in sub-Saharan Africa face considerable challenges with WASH. While 1 in 4 healthcare facilities do not have basic water services (water from an improved source in the hospital premises) globally, this figure stands at 51% for sub-Saharan African facilities. Data provided from 28 countries shows 23% of facilities in this same region have improved sanitation facilities(separate toilets by sex, toilets for those with limited mobility and separate toilets for hospital staff). Healthcare waste management is also highlighted as a challenge in many of these African countries with only 40% of facilities safely segregating healthcare waste(7).

There have been renewed efforts to improve WASH services with the WHO and UNICEF and partners all over the world working to *“achieve universal access to WASH in all facilities in all settings by 2030”* (4). As part of this initiative, core and extended indicators to track WASH in HCFs were developed, tested and revised, a research agenda was drawn to address the evidence gaps and the “Water and Sanitation for Health Facility Improvement Tool” (WASH FIT) was developed alongside training material for facilities in resource-poor settings to use (8).

WASH FIT was not designed for national or regional level situation analysis, monitoring or tracking of WASH in hospitals. Instead, the tool guides teams of staff and community representatives within small primary facilities (e.g. those providing outpatient services, family planning, antenatal care and uncomplicated delivery care) through a continuous cycle of assessing and prioritizing risks linked to poor WASH, defining and implementing improvements and continually monitoring progress locally and autonomously. WASH FIT focuses on actions involving maintenance and repair as well as infrastructural and behavioral change, which are ideally integrated into broader quality improvement plans. WASH FIT covers four broad domains (Fig 1) and comprises 65 indicators and targets for achieving minimum standards for maintaining a safe and clean environment. Minimum standards are based on global standards as set out in the WHO essential environmental health standards in healthcare(9) and the WHO Guidelines on core components of infection prevention and control (IPC) programmes at the national and acute healthcare facility level (10).

**Fig 1.**
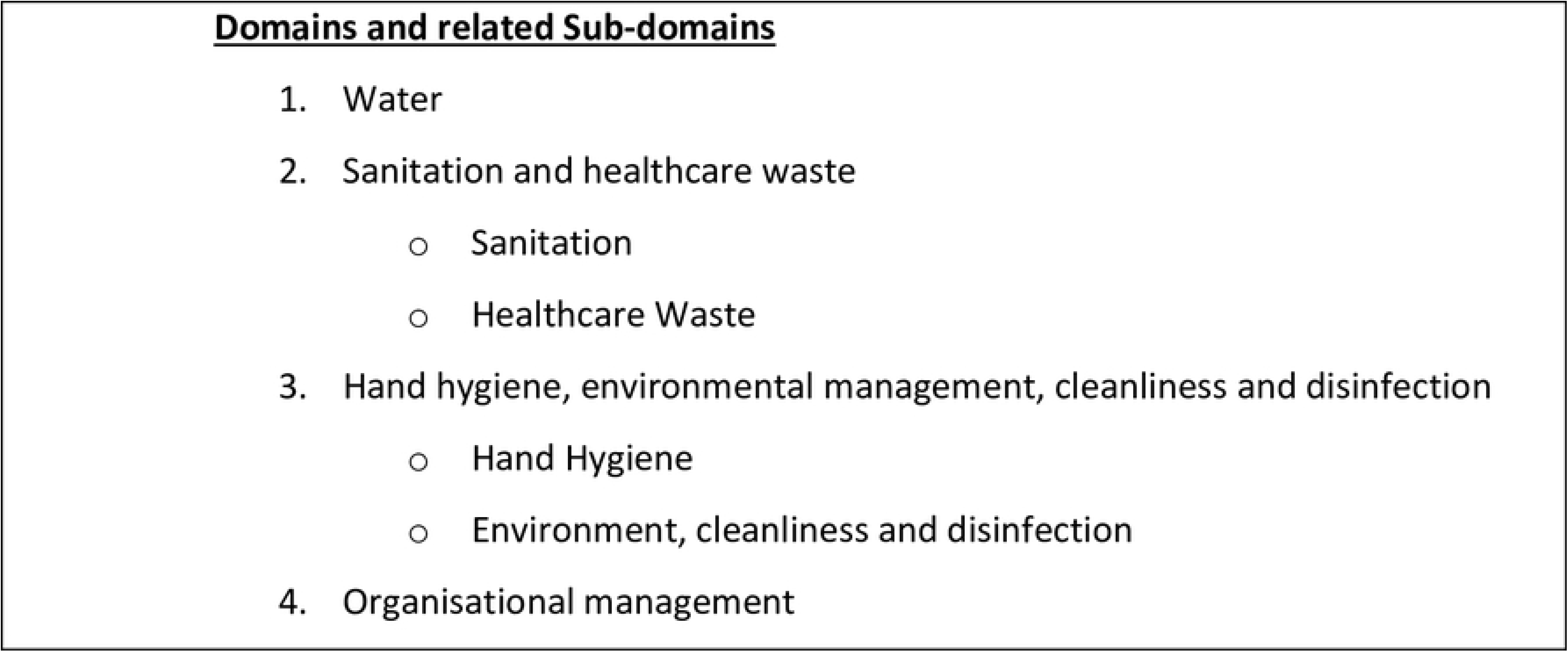
Domains assessed in WASH FIT

WASH FIT involves scoring 65 indicators using a three-level qualitative system (meets, partially meets, or does not meet the required standard) but has not been used to generate an overall facility or domain scores. Larger facilities (e.g. referral hospitals) raise additional, specific issues. They have more complex management and leadership arrangements(8) and deliver both inpatient and outpatient care spread across multiple wards, departments and service areas.

WASH FIT focuses on local improvement rather than the broader health system context and capacity. In Kenya, the provision of health services is the responsibility of county governments. County (formerly district) hospitals are headed by a medical superintendent/director who reports to the county health department. The hospital health management team comprising the medical superintendent, health administrative officer, nursing officer in charge and the departmental heads are involved in the day-to-day running of the hospital (11). These teams are in turn assisted by different hospital committees constituted within the hospitals; these include the IPC committees. The hospital managers and committees prepare budgets and staffing needs but the final budgetary and human resource allocation to these hospitals is the prerogative of county government (11). Larger hospitals in many low- and middle-income countries (LMICs) have similar organisational arrangements and some similar form of regional administration which have a role in decision making and resource allocation. These need to be involved in quality and WASH improvement efforts.

Our report describes an adaptation of WASH FIT which includes an extension of the tool to meet both local, national and regional needs for improvement and tracking linked to comprehensive assessment of WASH services in hospitals providing both outpatient and inpatient care. The adapted tool, WASH FAST (Facility Survey Tool) also considers how responsibilities should be allocated across different levels of the health system to promote accountability and improvement.

## METHODS

### Adaptation of WASH FIT to WASH FAST

The adaptation of WASH FIT to WASH FAST entailed four main steps: (1) developing a numeric scoring system and a system for meaningful aggregation of scores, to enable comparison across HCFs and facilitate tracking of WASH services over time; (2) modifying the facility assessment – so that indicators are assessed and scored for each ward in addition to the facility as a whole – to identify potential variation in WASH across HCF service areas and; (3) identifying those responsible for effecting positive change in WASH. The fourth step involved demonstrating feasibility and potential value of the WASH FAST.

#### 1. Development of a numeric scoring system

The first step involved developing a simple quantitative scoring system that assigns a numeric score to each indicator based on assessment findings as follows: 0-does not meet the required standards (i.e. target), 1-partially meets target and 2-fully meets target. This enabled us to create aggregate domain scores (based on the number of indicators within a domain) and aggregate facility scores (based on all 65 indicators) that can be used to compare domain and facility performance using both a numeric and “traffic light” reporting approach.

#### 2. Identification of ward level indicators

The second step involved identifying and adapting certain indicators that can independently and objectively be assessed at the inpatient-ward level. This was an iterative process where the research team (19 health professionals comprising doctors, nurses, pharmacists and public health officers who had been recruited to apply the WASH FAST assessments in hospitals in Kenya) reviewed all 65 indicators and identified which were suitable for ward-level assessment, then tested the indicators to finalize the tool. Using the same simple numeric scoring approach as in Step 1 above enabled exploration of variation in performance between wards in the same hospital.

#### 3. Assigning responsibility for action

The third part of the adaptation was to create new domains of indicators based on who should take responsibility for their action – addressing the issue of accountability within and beyond the facility. For this process a study team of four members examined all 65 indicators aiming to understand how these indicators are related while also assigning each indicator to the person(s) responsible. The levels of responsibility were validated through a series of interviews with healthcare workers and a subsequent stakeholder workshop.

#### 4. Demonstrating potential use and creating tools to help visualise variation in performance

We proceeded to collect data using the WASH FAST tool as part of a survey in 14 county hospitals, during which key informant interviews were also conducted. The county hospitals vary in size and are from 11 counties in both high and low malaria zones in Kenya (5 and 9 sites respectively). Hospitals were purposefully selected based on links developed from ongoing work to improve clinical information as part of a collaboration between the KEMRI-Wellcome Trust Research Programme and the Ministry of Health (MOH) (12).

WASH FIT entails five main tasks as part of a continuous improvement cycle, namely; (1) assembling a team; (2) conducting a facility assessment using a qualitative scoring system for each domain indicator; (3) undertaking a risk and hazard assessment; (4) developing an improvement plan; and (5) undertaking a continuous evaluation process. This study only carried out the first two steps, assembling a team and facility assessment. At each hospital, teams consisted of 7-8 people. These were a study team leader, 4 surveyors employed for the study and 2-3 representatives from the individual hospitals where they survey was being carried out, selected based on their specific role as IPC coordinators or public health officers. Data collection involved direct observation and hospital workers provided clarification of the assessment where needed. Each indicator was assessed, and the score (not meeting target, partially or fully meeting the target) determined by team consensus. Data were collected for each inpatient ward (34 indicators) then for indicators assessed at the whole facility level (65 indicators). The 65 facility level indicators included an assessment of outpatient areas, common service areas (e.g. kitchen, laundry, laboratory, waste management facilities) and the outdoor environment. The facility level assessment also included the 34 indicators assessed at ward level. For these indicators scoring took account of all ward-specific scores together with all other relevant hospital areas and was based on the overall judgement of the survey team. The data collection tool and the standard operating procedures are provided as an annex.

Aggregate scores were generated by summing individual indicator scores and dividing this total by a denominator that assumed a perfect score for each indicator. In this way we then estimated percentage scores for the hospital, WASH domain and level of accountability. These summary scores were based on the facility level indicators for the whole hospital or indicators relevant to each WASH domain or level of accountability respectively. Summary ward-specific scores were based on individual indicator assessments made for each ward. The rationale for such sub-scores is to enable exploration of performance variation to help identify priority areas for improvement and assign responsibility for improvement. To enable interpretation of scores, we generated a revised ‘traffic-light’ colour maps presenting percentage scores using cut-offs of <40%, 41-60%, 61-80% and 81-100%. Data analysis for visualisation was done using R, an open source statistical package (13).

Qualitative data collection included 31 ‘long’ or ‘ethnographic’, semi structured interviews conducted during the study period between February and April 2018. Seventeen interviews were conducted with people with a managerial role. These were medical superintendents, nursing officers in-charge, chief pharmacists, laboratory managers, departmental heads and ward managers. Fourteen interviews were with frontline healthcare workers including consultants, medical officers, nurses, laboratory technologists and pharmacists. These interviews served two purposes: primarily to understand IPC arrangements in Kenyan hospitals (with findings reported elsewhere), and to explore the likely validity of our proposed allocation of indicators for accountability. For the interview data, all interviews were recorded and transcribed verbatim and the transcriptions verified by the authors (GK and JM). They were then coded into themes providing key insights into IPC arrangements in Kenya. Interview data were also examined for evidence that confirmed or revised the levels of accountability assigned in other processes. Coding of interview data was performed using NVivo version 12.

To validate and demonstrate the potential for this tool, we undertook two forms of validation: analysis of interviews with healthcare managers and staff for confirmation or revision of our ideas and then presentation and discussion of the tool at a stakeholder consultative workshop. This workshop included 120 experts and key stakeholders in WASH comprising: Ministry of Health officials, hospital WASH leaders, county health department leaders, and doctors and nurses in Kenya with an interest in IPC. The workshop aimed to establish face validity for the levels of responsibility assigned to each indicator and the value of presenting data using aggregate scores.

## RESULTS

We established that 34 of the 65 indicators could be assessed independently and objectively at the ward level. A description of all 65 indicators is provided as a supplement. Table 1 below provides a summary of the number of indicators that were to be assessed at the ward and facility level by the original WASH FIT domains and by the proposed levels of responsibility.

### Responsibility for action

For the process of assigning responsibility for action, the research team developed a provisional reorganisation of indicators based on their logical relationship and who it was felt likely would be responsible for action. We classified three levels of responsibility. These are, first, the county government which should be concerned with indicators that are beyond the control of hospital leadership. The second level is the hospital health management team (the medical superintendent, health administrative officer, nursing officer in charge and the departmental heads) and thirdly, the hospital infection prevention and control committee. This is as shown in the Table 1 below.

Two of the 9 indicators under the responsibility of the county government could also be assessed at ward level ((i) water services available in sufficient amounts and (ii) having rewards for high performing staff), these only need to be assessed at the facility level for tracking progress in follow up assessments. Therefore, when grouped by responsibility we suggest only 32 of the original 34 indicators are assessed at ward level as shown in the Table 1.

**Table 1:** Summary Indicators at ward and facility level by WASH domains and WASH FAST

| WASH FIT |  | WASH FAST |  |  |  |  |  |
| --- | --- | --- | --- | --- | --- | --- | --- |
| WASH DOMAINS |  | WASH DOMAINS |  |  | ACCOUNTABILITY DOMAINS |  |  |
|  | Facility |  | Ward | Facility |  | Ward | Facility |
| Water | 14 | Water | 6 | 14 | County Government | 0 | 9 |
| Sanitation & Healthcare Waste | 22 | Sanitation & Healthcare Waste | 11 | 22 | Hospital Management | 16 | 31 |
| Hand hygiene, Environmental Management, Cleanliness and Disinfection | 18 | Hand hygiene, Environmental Management, Cleanliness and Disinfection | 12 | 18 | Infection prevention & control committee | 16 | 25 |
| Organisational Management | 11 | Organisational Management | 5 | 11 |  |  |  |
| **Total** | **65** | **Total** | **34** | **65** | **Total** | **32** | **65** |

We then examined interview data to see whether the initial re-organisation was valid or needed modification.The in-depth interviews allowed us to explore the relationships between the WASH criteria and to establish where responsibilities lay for each. This was an important step, because it contributed to the emerging model of the layers of WASH management and informed our understanding of the causalities and contingencies in this area.

County Level – The County is responsible for setting the budget for each hospital, and importantly, sets the overall budget for health spending. This impacts on hugely important, WASH-related criteria such as staffing levels and material upkeep of hospitals. Additionally, while each department in each hospital is asked to project their needs for the next year as part of hospital budgeting processes, the requested amounts may be ignored by counties. Hospitals thus need to work within the limitations of the budget and staffing allowed them.

> *“You know normally we are told to itemize whatever we require in the departments that we are working in. …yes, by different departments come up with their budget proposal. The administrator compiles the budget for the whole hospital and then gives it…we don’t control funds in the institution. Every finance that is channelled to the hospital is controlled by the chief officer in the county. So, we send the budget to the county” <u>Hospital Manager</u>*

Whereas the day-to-day running of the hospital is done by hospital management, some of the activities are delegated to committees within the hospital.

Hospital Level – Key areas of hospital management were delegated to committees that held responsibilities for activities and addressing needs. The effectiveness of IPC committees appeared to be variable between hospitals, but where they were operational, they had an important influence on resource allocation and monitoring of WASH.

> *“…when the committee, the IPC committee, meets they raise their needs as per various departments, and then the hospital now addresses that. Like if you want to purchase, for instance…bins, litter bins, disposal bags, waste disposal bags. So, you just raise your needs as per your department because you know different departments have got their different needs” <u>Hospital Manager</u>*

However, despite their importance, these IPC committees in some facilities struggled to gain respect relative to other more prestigious committees e.g. the procurement committees and were regarded to be of low status.

> *There are some committees which are found to be more, which are more do I say prestigious? They look better. So, if I am in IPC, people will be thinking okay… so IPC will have no one. I mean what is the benefit of being in IPC, what is there, how am I gaining being in IPC? <u>Consultant</u>*

Ward Level – Interest and capacity at the ward level is essential for providing effective WASH services. The individuals responsible for WASH at this level are not likely to have the ability to affect budgets and resource allocation but they are important in both maintaining supplies and overseeing important areas such as hand hygiene. And more broadly in championing and advocating for the importance of WASH. Variability in performance at ward level may be linked to the presence of an individual in the ward who has interest and passion for IPC-related activities.

> *“And we also have someone, he’s also a team leader in the infection control and making sure we have… whenever he’s available we have our sanitizers, make sure we have soap, make sure we have gloves” <u>Frontline Health Worker</u>*

The relative importance of IPC varied ward to ward, however, with the newborn units (NBU) repeatedly cited as an example to compare high versus low performance:

> *“Across the hospital, in NBU is where I know there is strict infection prevention because once you are getting into NBU, you remove your lab coat, you wash your hands and then you get into the unit where you fold your, whatever if you are wearing a long sleeved anything you fold it and then you get in the unit… Now in other wards we don’t have such strict infection prevention, you just get in and you start…”* <u>Frontline Health Worker</u>

### Consultative Workshop

An updated arrangement of indicators linked to the three levels of responsibility, was presented to stakeholders. The consultative workshop was conducted in November 2018 during the annual National Infection Prevention and Control Symposium. There were 120 people in attendance at the consultative workshop. These included Ministry of Health officials, managers from the hospitals and county government, development partners and training institutions who are familiar with IPC and WASH, and frontline health workers. The workshop attendees largely agreed to the proposed levels of accountability with key recommendations for hospitals to identify a champion to lead the IPC committees and to identify ways of boosting morale for IPC-related issues among health workers across these hospitals. This resulted in the final framework (Fig2) which shows the relationship between indicators, their original domains, new levels of responsibility and highlights which individual indicators can also be assessed at ward level. From the figure we note the roles played by the WASH/IPC focal persons who are members of the IPC committees in conducting training, ensuring that there are WASH materials and assisting in conducting audits to assess availability of WASH materials around the hospital.

**Fig 2.**
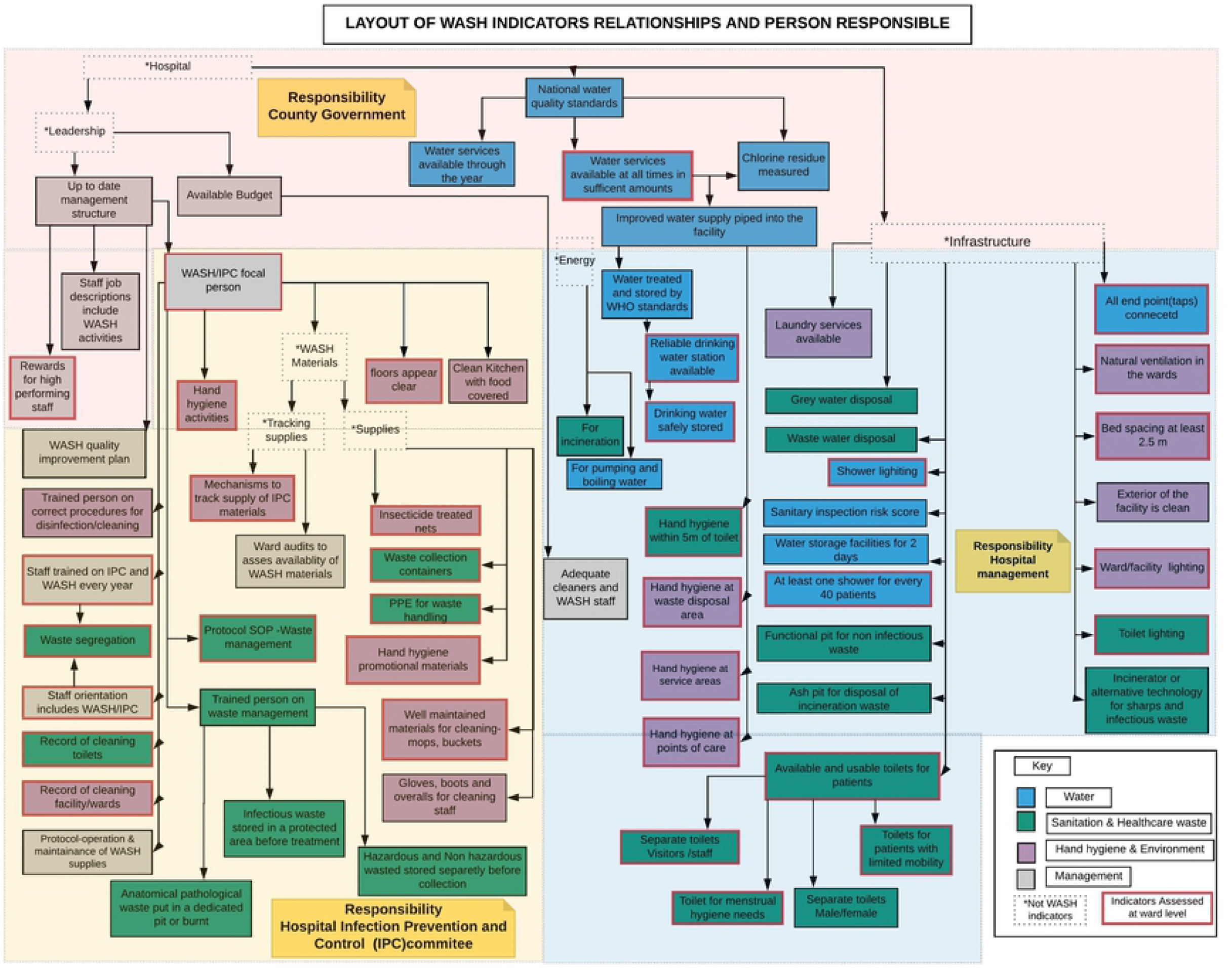
Schematic layout of WASH FIT indicators.

These shows how indicators assessed at ward and facility level logically related. These are grouped by the original 4 domains and by levels of responsibility. The indicators with a red bold outline were also assessed at ward level. The dotted boxes are used to describe categories and are not part of the indicators.

#### Visualization approaches to support monitoring

Using an example of data collected from four of the 14 hospitals, two large (H2, H9) and two small (H1, H7) hospitals, we present an illustration (Fig 3) of how performance of two domains (water and sanitation) varies between hospitals (Panel A) and how the individual wards within these facilities performed (Panel B). For the two domains we note marked variability across the hospitals with some facilities having scores of <50%. We also note variability between and within wards in these hospitals. From this example (Panel B), the water domain ward scores in hospital H1 show minimal variability compared to those of hospital H9. The figure also presents data from the original WASH FIT at facility level for the four hospitals presented (Panel C)

**Fig 3.**
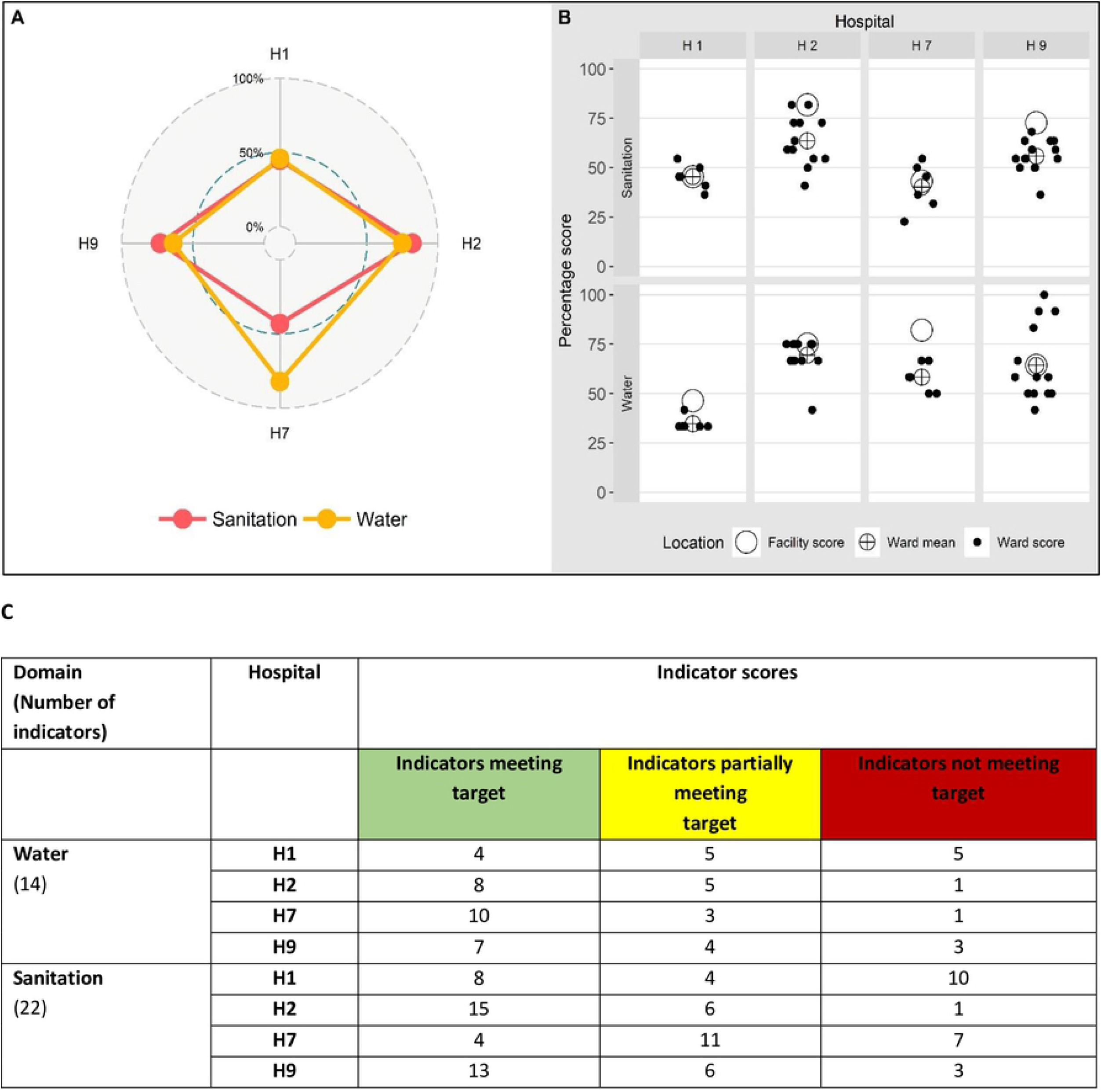
Service performance variation by ward and hospital and the original WASH FIT scores. (A): Radar Plot of facility level scores from 4 hospitals for the 2 domains (Water, Sanitation) showing similar performance for sanitation overall but more marked variation for water varying hospital performance. (B)Shows ward domain scores from multiple wards (dots) for 2 domains (water, sanitation) illustrating their variation, the mean of these ward-specific scores (⊕) and the overall facility aggregate score(O). The overall facility score includes assessment of inpatient wards and other service areas (kitchen, outpatient, outdoor environment) across the hospital. (C) shows WASH FIT facility level scores of 5 hospitals for the 2 domains of Water and Sanitation.

To further illustrate how the approach can provide detailed information for use at national and regional levels on hospital performance and who needs to act, we present an example of the 16 indicators [spanning all the WASH domains] which are the responsibility of the IPC committee at ward level. Here we generate the summary ward scores for each of the four hospitals (H1, H2, H7, H9) coded using a traffic light colour system with red being a score of <40% and green indicating a score of >81%. Fig 4 provides an illustration of a ‘dashboard’ approach that shows performance across the hospitals for each IPC committee-related indicators assessed at ward level (horizontal bar chart, for example highlighting a need for the IPC committees to work on availing cleaning records in the wards in these 4 hospitals). It also helps visualise the overall mean ward scores for each hospital for all 16 indicators (the top panel vertical bar chart) and the traffic light coding presents a summary of how the individual indicators performed in individual hospitals.

**Fig 4.**
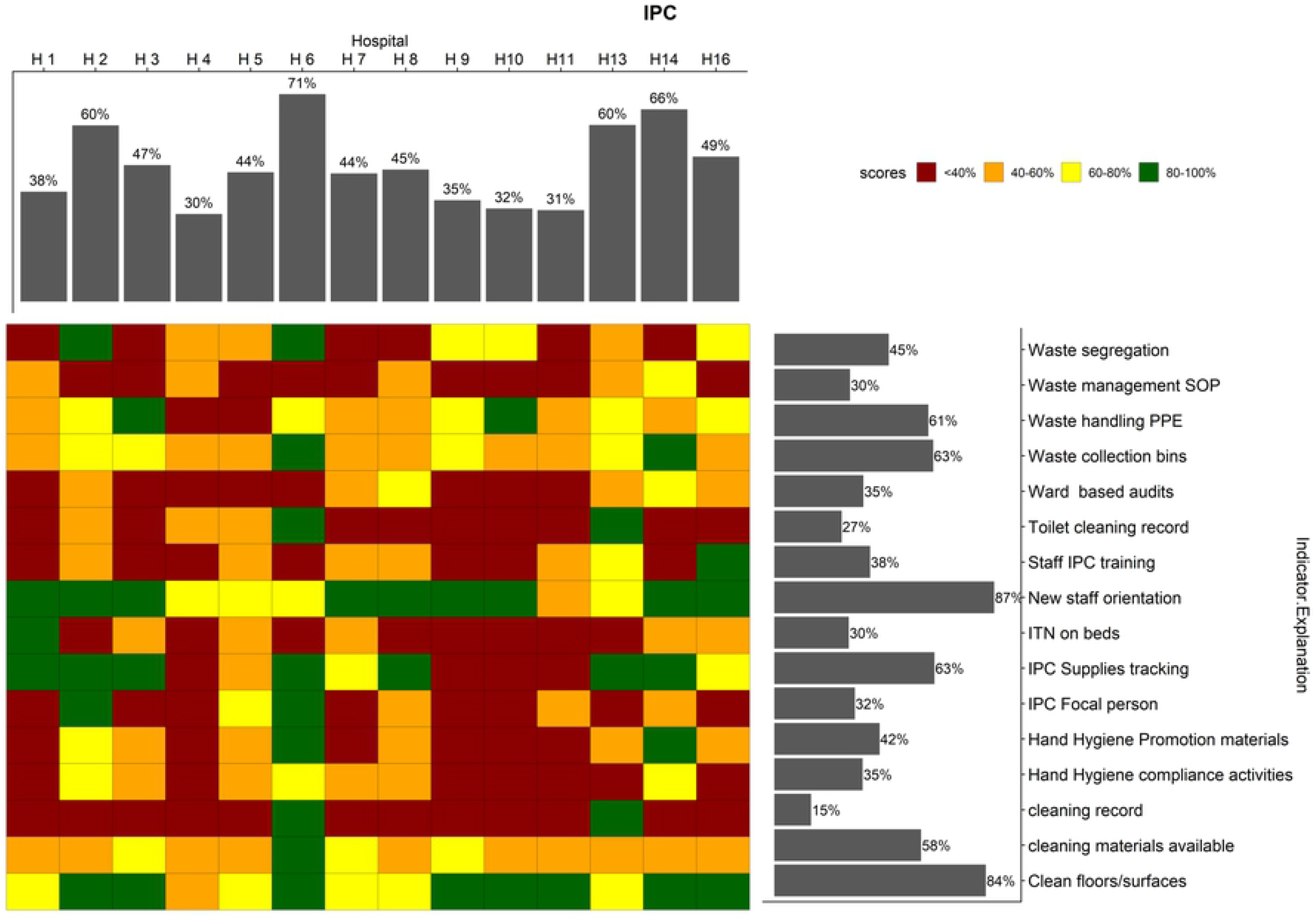
Performance of Infection prevention and control domain indicators. The mean service performance at ward level for the indicators under the IPC committee is shown by the upper bars. The right bars summarize the performance of each indicator across the 4 hospitals. The squares in the central grid are coloured according to the performance classification of each indicator in each hospital, by the colour categories. SOP: standard operating procedures, PPE: personal protective equipment; IPC: infection prevention and control; ITN: insecticide treated nets

## DISCUSSION

We have presented an adaptation of the Water and Sanitation for Health Facility Improvement Tool (WASH FIT) into WASH FAST (Facility Survey Tool). The adaptation entailed an extension of the tool to meet both local (i.e. self-improvement) and national (i.e. situation analysis, monitoring and tracking) needs, and to facilitate comprehensive assessment of WASH services in larger – more complex – secondary and tertiary facilities providing both outpatient and inpatient care and multiple medical specialties. The adapted tool scores indicators at various levels of the facility (including by ward and by medical specialty) and assigns levels of accountability for each indicator, to identify what services can be addressed by who locally or at higher levels of the health system. An aggregated numeric scoring system, consisting of a percentage score out of the total that would be obtained if all indicators met the expected target, can be used to locally identify service areas requiring priority action within a facility, or to identify facilities or specialties requiring priority action nationally or sub-nationally.

Adequate governance and leadership are one of the foundations for provision of quality care. Governance for quality includes improving accountability and identifying the roles and responsibilities at all levels of the health systems and using data to make decisions (14). Thus WASH FAST may also be used to identify responsible actors limiting or effecting positive change within a facility and/or beyond, and to potentially reward good performance in a bid to improve staff morale in regards to IPC/WASH. WASH FAST assumes that performance and quality indicators for WASH are the responsibility of three possible actors, namely administrative division officers or governments responsible for budgetary and human resource allocation to hospitals, senior hospital management teams and relevant facility-based specialised committees or groups of persons who are key in decision making for IPC-related activities, such as the infection prevention and control committee. Although the WASH FAST was generated within a Kenyan context, we expect these broad accountability domains (validated by government representatives, public health officers, IPC experts and healthcare professionals through interviews and a consultative workshop), to be generally applicable to most LMICs, with minor adaptations to be made according to the context, to allow for within-or between – country comparisons where applicable.

We anticipate that aggregated scores derived from application of WASH FAST can be used more broadly to inform health system leaders on what actions facilities and specialties require at either local, sub-national or national level to improve WASH services. The numeric system allows the comparison of WASH services within- and between-facilities and or medical specialties either cross-sectionally or over time (i.e. to identify changes in quality and performance and trends), through repeated surveys. The extended tool is hence a broadly applicable facility improvement tool – potentially encompassing training, team-building and risk assessment steps as per the WASH FIT process – that also appertains to WASH performance monitoring sub-nationally and /or nationally. Training of facility staff to partake in surveys and/or facility improvement plans, in turn empowers and encourages staff to take interest and ownership on WASH and IPC, contributes to up-skilling in these areas and improves short and long term sustainability of interventions and developments where applicable (15). WASH FAST may also be applied to help remedy the paucity of data on the status of WASH services in LMICs, help bridge evidence-based gaps and provide a platform to monitor interventions aimed at improving WASH and patient safety. This is whilst continuing to serve the original purpose of continuously informing a local improvement plan in small primary HCFs as well as larger facilities comprising multiple wards and medical departments.

A limitation of both WASH FAST (and WASH FIT), is that the score assigned to selected individual indicators may be subjective, where it relies on observations that could vary from person to person. To mitigate this, we developed standard operating procedures before data collection, conducted training for the data collection teams and used consensus among surveyors to assign scores during the assessments. WASH FAST also rests on the premise that the hospitals have well-structured leadership including a functioning infection prevention and control committee or relevant expert group. The indicators are also not weighted in accordance to the health risk they pose, implying that identical aggregate scores may have very different decision-making implications depending on the composite of indicators considered in the score (e.g. availability of water vs. cleaning protocols). The same principle applies to overtime tracking, where a facility may get the same score over two consecutive surveys, perhaps reflecting improvements in some areas but worsening in others. To mitigate these limitations, aggregated (summary) scores should be interpreted in the context of individual indicator scorings presented through heatmaps or other visualisation tools.

In a bid to improve hospitals as platforms that provide high quality care, proper WASH and IPC structures are at the core of this provision and cut across most components for the high-quality health system as recommended by WHO (14). We suggest that using WASH FAST to monitor and improve the capacity for WASH and IPC, would also improve governance for quality which includes improving accountability and identifying the roles and responsibilities at all levels of the health systems (14). In the process of accelerating universal health coverage in many counties, hospital accreditation has become a key component as it provides a criterion for insurers and governments to include in their funding mechanisms. WASH FAST can thus be used as part of the tools for hospital accreditation activities as part of these assessments include the hospitals IPC structures and can also provide data to the District Health Information System (DHIS) (16) (17).

## CONCLUSION

We propose the WASH FAST (Survey) tool developed can support situation analyses, performance tracking and effective decision making for WASH and IPC at different levels of the health system for larger facilities such as hospitals in resource limited settings. Where the primary aim is local improvement in smaller facilities WASH FIT remains the tool of choice.

## ACKNOWLEDGEMENTS

We would like to thank the Kenyan Ministry of Health and the Council of Governors who gave permission for this work to be carried out. We also thank the hospital management and clinical teams who supported the work in the survey hospitals. This work is published with the permission of the Director of KEMRI.

## AUTHOR CONTRUBITIONS

The roles of the contributors were as follows: M.Maina, J.M, O.T, C.S and M.E conceived the study. M.Maina,G.K,M.Z,O.T collected data, P.M, O,T,J.M assisted M.Maina in analysis and interpretation of these data. M.Maina, AH,M.M, O.T, C.S and M.E drafted the manuscript. M.Maina,A.H,J.M,O.T, C.S and M.E critically revised the manuscript for intellectual content. All authors read and approved the final manuscript.

## AVAILABILITY OF DATA AND MATERIAL

All summary data underlying the findings is freely available in the manuscript and supplemental files, however since this was data collected in collaboration with the Ministry of Health and under terms of ethical approval granted by KEMRI (SSC Number 3450) and the Ministry of Health. The existing ethical approval and agreements with the Ministry of Health do not provide for the data set to be hosted in a public repository. Access to these raw data may require additional approval from the Ministry of Health and submission of a proposal for ethical review. Requests can be facilitated by contacting the corresponding author.

